# A proposed rule for updating of the head direction cell reference frame following rotations in three dimensions

**DOI:** 10.1101/043711

**Authors:** Jonathan Wilson, Hector Page, Kate J. Jeffery

**Author notes:** Kate J. Jeffery, Institute of Behavioural Neuroscience, Department of Experimental Psychology, Division of Psychology and Language Sciences, University College London WC1H 0AP, UK.

## Abstract

In the mammalian brain, allocentric (Earth-referenced) heading direction, called azimuth, is encoded by head direction (HD) cells, which fire according to the facing direction of the rat’s head. If the animal is on a horizontal surface then egocentric (self-referenced) rotations of the head around the dorso-ventral axis, called yaw, correspond to changes in azimuth, and elicit appropriate updating of the HD signal. However, if the surface is sloping steeply then yaw rotations no longer map linearly to changes in azimuth. The brain could avoid the complex computations needed to compute global azimuth simply by always firing according to direction on the local surface; however, if the animal moves between surfaces having different compass orientations then errors would accumulate in the subsequent azimuth signal. These errors could be avoided if the HD system instead combines two updating rules: yaw rotations around the D-V axis *and* rotations of the D-V axis around the gravity-defined vertical axis. We show here that when rats move between vertical walls of different orientations then HD cells indeed rotate their activity by an amount corresponding to the amount of vertical-axis rotation. With modelling, we then show how this reference-frame rotation, which may be driven by inputs from the vestibular nuclei or vestibulocerebellum, allows animals to maintain a simple yaw-based updating rule while on a given plane, but also to avoid accumulation of heading errors when moving between planes, thus facilitating orientation during complex real-world navigation.

## Introduction

The neural circuitry supporting navigation has been intensely studied for several decades, and attention is now turning to how navigation processes operate in the complex, three-dimensional real world. We consider here the problem of maintaining a stable heading direction signal when moving over a non-flat surface.

Computation of head direction, or azimuth, is a core feature of navigation, and in mammals this faculty is supported by the head direction (HD) neurons, the firing of which is azimuth-sensitive (Fig. 1A). In a horizontal environment, HD neurons are updated by changes in head direction occurring as the animal rotates around its D-V axis: these rotations are called *yaw* (see Supp. Fig. 1 for the complete set of rotational axes and terminology). During yaw, a simple transfer function moves activity through the HD cell network, often conceptualized as a ring attractor(Skaggs et al. 1995; Zhang 1996; Redish et al. 1996); Fig. 1A) so that their activity always reflects the animal’s current azimuth: yaw thus maps linearly to azimuth change.

**Figure 1.**
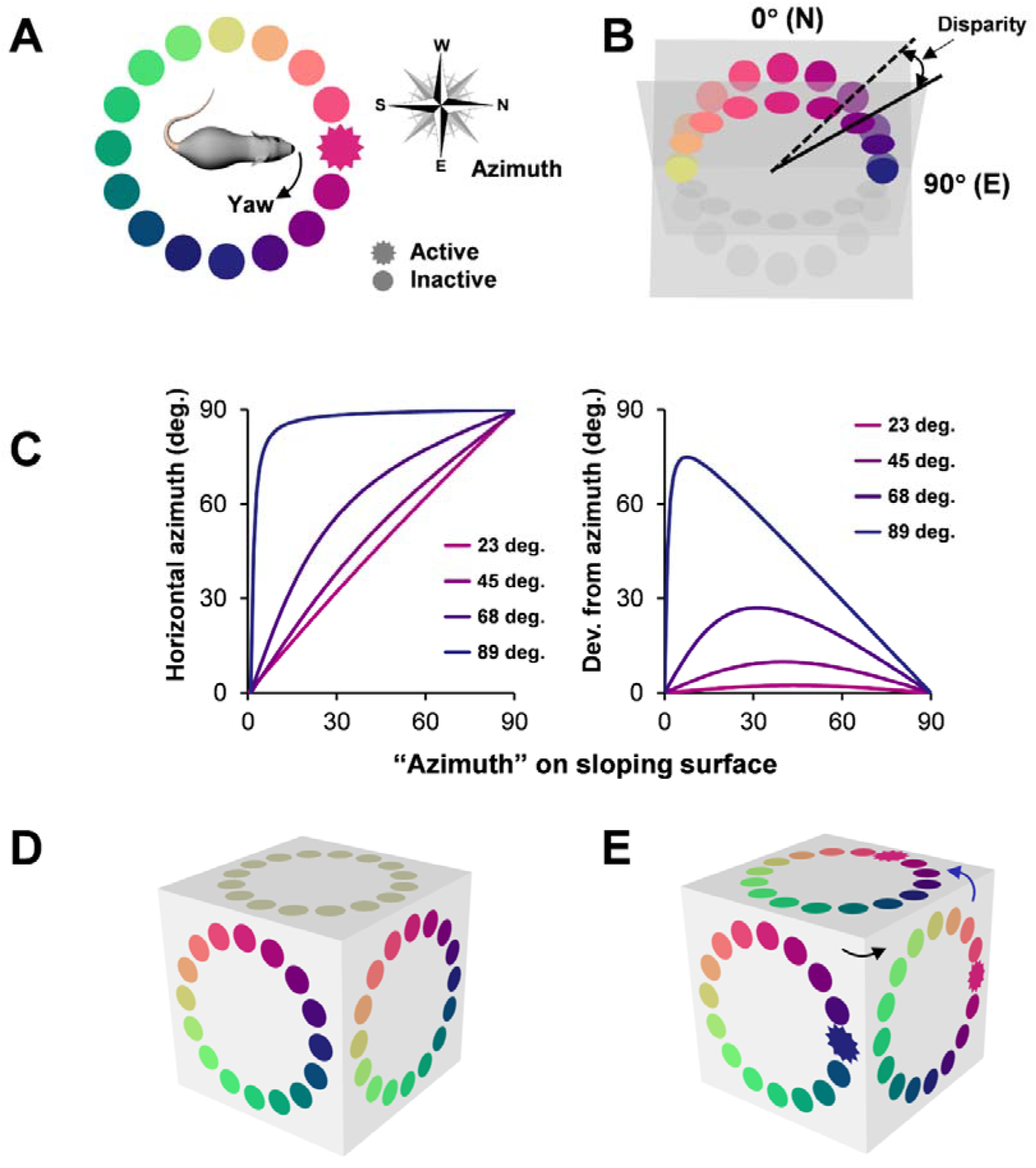
(A) Schematic of the conceptually ring-like organization of head direction cells in which different cells encode different directions (“azimuths”). The active cell represents the direction the rat is currently facing, and a head turn (“yaw”) moves activity around the ring. (B) Slope-dependent mismatch in mapping between yaw angle on a sloping surface and azimuth on the horizontal plane. Colored circles represent head direction neurons, operating either on a horizontal plane or on a plane tilted up towards the viewer by 60 degrees. Note that on the tilted plane, the firing direction of many neurons have changed their azimuths. While zero and 90 degrees neurons have unaltered azimuths, there is a progressively increasing change in the intermediate directions. (C) Quantification of the effect shown in B: Graphs showing the relative azimuths on horizontal and sloping surfaces (left) and the disparity between these values (right) as a function of surface slope (legend), and of compass direction (x-axis). The mismatch between sloping and horizontal azimuths is slope-dependent, becoming increasingly non-linear as the slope steepens. A network trying to encode azimuth would thus have to change the relationship between yaw and azimuth in a slope-dependent manner. (D) Inconsistency between orientations of the HD network on three orthogonally abutting surfaces, occurring when the patterns on the vertical surfaces are locally specified (up/down remain up/down, left/right remain left/right, etc). On the top surface, there is no arrangement of the colored neurons that is consistent with both of the abutting vertical surfaces at the same time. (E) The inconsistency problem can be solved using a rule that rotates the network accordingly when the rat turns a vertical corner (black arrow) but not if it rounds a horizontal one (blue arrow). For this left turn, activity (starred unit) moves to the left around the ring, because its reference frame has rotated to the right. Note that there is now a configuration on the top surface that is consistent with both of the others.

Computing a stable azimuth signal during movement over a non-flat surface, however, poses challenges. Two important problems are (1) slope-dependent non-linearities in the transfer function that shifts activity through the network from one set of cells to the next as the animal turns its head (Fig. 1B and C), and (2) conflicts in azimuth computation at a given location following movement over differently-oriented non-horizontal surfaces (Fig. 1 D and E). Here, we provide evidence that such non-linearities and conflicts can be avoided by updating the head direction signal not just by rotations of the head about the dorso-ventral (D-V) axis, which is an egocentric (self-centered) reference, but also by rotations of this axis around the gravity-defined vertical axis, which is an allocentric (earth-centered) reference. In this way, the HD cells can remain consistently oriented in allocentric space, thus allowing for orientation even during navigation over complex terrain.

We examined the case of an animal moving over differently oriented vertical surfaces, and formulated three alternative hypothetical encoding schemes to predict how HD cells would behave (Fig. 2) – “local,” in which the cells treat each surface in isolation and update only according to head rotations on that surface (called *yaw* rotations); “global,” in which the cells respond to rotations in any dimension in 3D space, and “mosaic” in which the cells update by a local rule at each given point on the surface *but also* if the D-V axis of the animal rotates around the vertical axis as defined by gravity. The mosaic rule would mean that as an animal rounds a corner from, say, a South wall to an East one, its D-V axis would rotate 90 degrees counterclockwise (seen from above) and the reference frame of HD cell activity would rotate 90 degrees clockwise (Fig. 1E), with the result that the actual activity within the ring would shift counterclockwise. Note that this is true irrespective of how the animal turns the corner (e.g., whether via roll or pitch).

**Figure 2.**
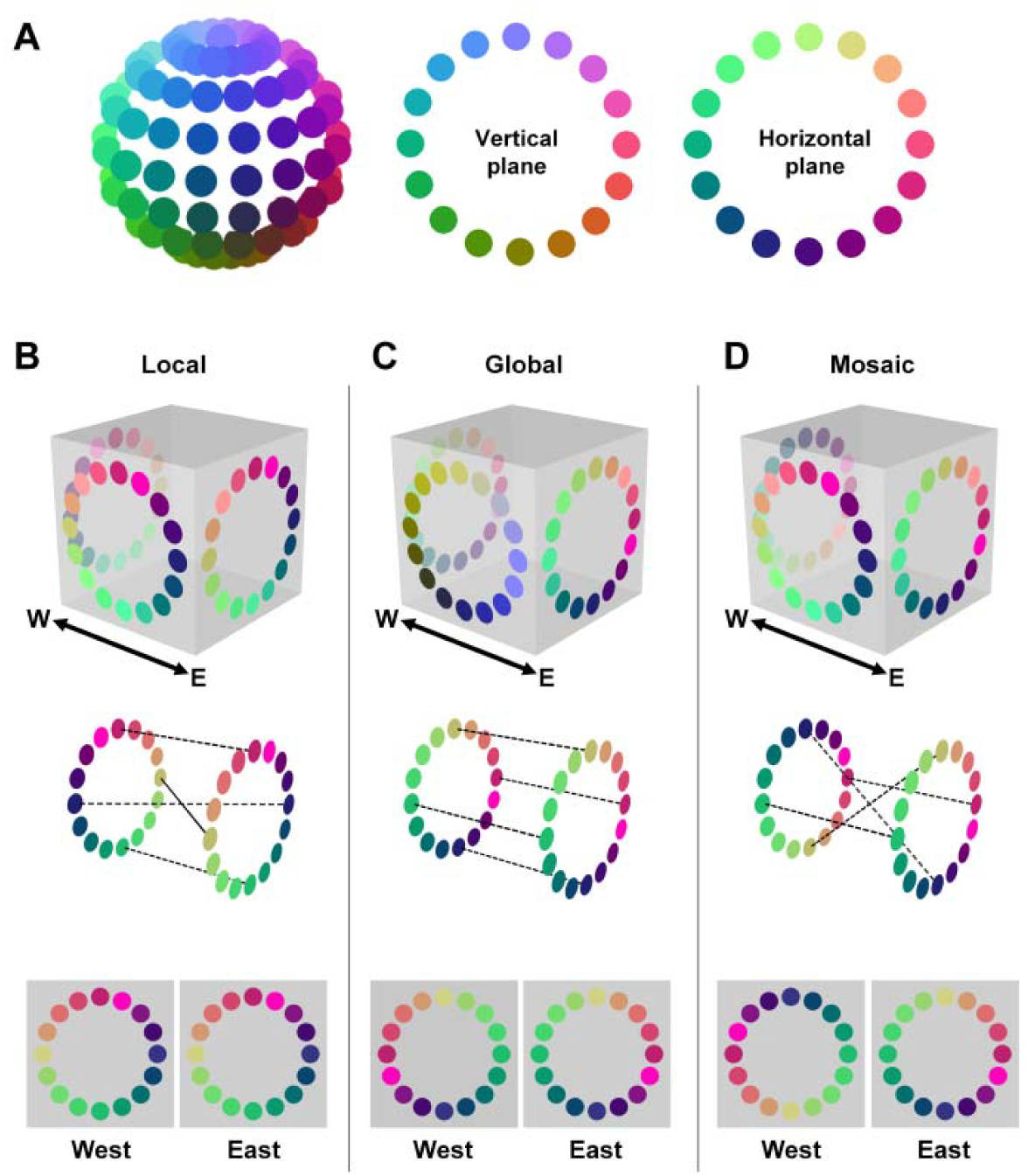
Hypothetical organizations of head direction cell activity in three dimensions. (A) The head direction cell attractor network might be configured as a sphere instead of a ring. Note that different cells would be active for slices taken horizontally vs. vertically through the sphere, as would be expressed on a floor vs. a wall. (B-D) Predictions made by alternative head direction cell coding schemes for a rat on a cuboid surface. The apparatus is shown as a transparent cube to allow visualization of the relative firing pattern on opposing surfaces. In a local scheme (A), the network is oriented purely by local egocentric cues – the same cell fires for “up” on all the vertical faces of the cube, and likewise for “down”, “left”, “right” etc. Note that firing on the top surface could not be consistent with all of the abutting vertical surfaces at once (see also Fig. 1D). (C) In a global scheme, the attractor is spherical like the one in (A): opposing faces show firing patterns that are spatial mirror images of each other when viewed facing each surface, meaning that rightward head turns need to move activity in opposite directions around the network on the opposing faces (from yellow to red to blue on the East wall and from yellow to green to blue on the West wall). On an orthogonal surface (the South face in this example), firing essentially reflects orthogonal slices through a spherical attractor and so a different subset of neurons would be active except for “up” and “down” cells, as illustrated also in (A). Yet another subset of neurons would be active on the top surface. In a mosaic scheme, the firing on each face is implemented by a simple ring attractor, but the orientation of the attractor is rotated between adjacent walls: to the right when the rat goes around the right-hand corner from the South to East walls, and to the left when the rat turns the left-hand corner from South to West walls. Note that firing on the top surface could be simultaneously consistent with all the abutting vertical surfaces: the red cell would fire to North, blue to East, light green to South and yellow to West. Furthermore, rightward and leftward head turns would always move activity in the same direction around the ring attractor regardless of surface orientation. (B) Predicted relative firing patterns on opposing walls given the three schemes shown in A. In the local scheme, firing patterns would look identical; in the global scheme, they would be mirror images, and in the mosaic scheme (local but also accounting for rotation around the vertical axis) the patterns would be 180 degree rotations of each other.

To distinguish these models experimentally we recorded HD cells from the anterior dorsal thalamic nucleus (ADN) as rats moved directly between the vertical faces of a cuboid (three-dimensional rectangle) box, the “tree trunk maze” (Supp. Fig. 2), while foraging for food. Two cameras, facing the East and West walls respectively, were used to record the direction of cell firing.

The three schemes make three predictions for how HD cells should fire on opposing walls:

1. The local scheme (Fig.2B) predicts that within a global (room) frame of reference, firing on two opposing surfaces would each be a 180-degree rotation of the other, while the patterns as seen from the perspective of the two facing cameras would be identical.
2. The global scheme (Fig. 2C) predicts that on opposing walls, firing would maintain a consistent direction relative to 3D space, while the patterns as seen from the two facing cameras would be mirror images. This model was proposed, but then discounted, by Finkelstein et al as a possible way to describe the HD firing patterns seen in bats (Finkelstein et al. 2015).
3. The mosaic scheme (Fig. 2D) predicts that within the global reference frame firing would be flipped vertically, while the patterns seen from the camera views would be rotated (rather than mirrored) with respect to one another. This alternative resembles the local hypothesis in that it preserves the same relative firing angles between neurons, but differs in that it allows firing to maintain consistency with the global (room) reference frame too. Using this scheme, a rat could climb onto the roof from either East or West walls without inducing HD cell conflict (Fig. 1E). A similar scheme, the hemitorus model, was proposed by Stackman et al (Stackman et al. 2000) to account for the preservation of HD firing between a floor and a wall; the mosaic rule extends their model by accounting for movement from one non-horizontal surface to another, and generalizes to movement between any two non-inverted surfaces (the case of inversion is discussed later).

## Results

### Single-neuron recordings

Of the 19 HD cells recorded in anterior thalamus (Supp. Fig. 3) from three rats (14, 3 and 2 cells respectively; see Supp. Fig. 4 for the full set), identified by their directional firing in baseline trials in a horizontal arena, all were active on at least one wall and 15 were directionally modulated on both walls. The other 4 cells (cells 2,4,5 and 9) were directionally modulated on one of the two walls but were not active on the opposing wall (defined as a rate less than 20% of the baseline rate); this may be because the animals undersampled the direction the cells might be expected to fire, given the mosaic rule (i.e., downwards, a direction that the animals tended to avoid). The data below are reported as means +/‐ s.e.m.

**Figure 3.**
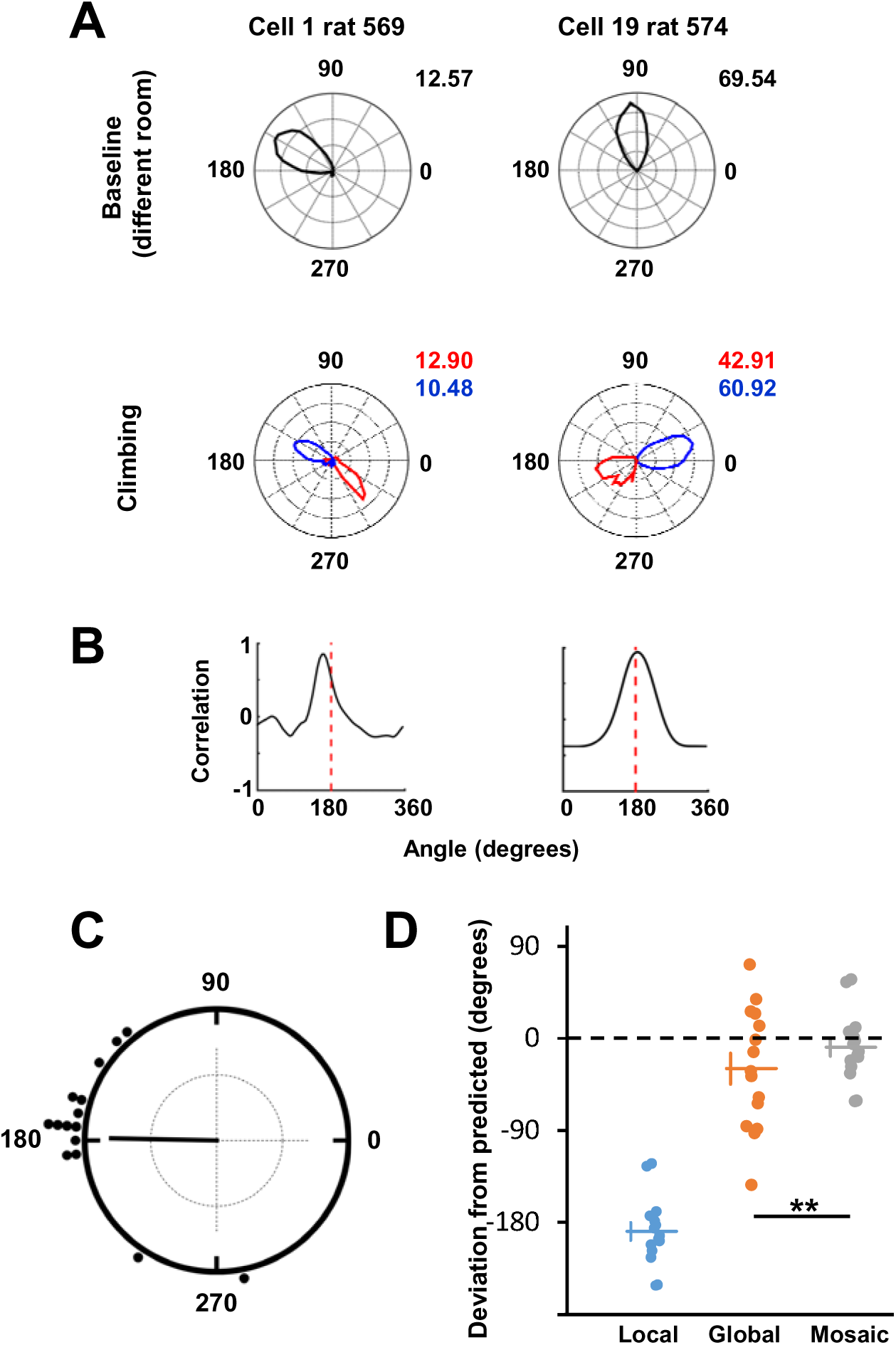
Properties of head direction cells recorded on the tree-trunk maze. (A) Polar plots of two cells, in baseline (top row) and tree-trunk (bottom row) trials, showing firing rate as a function of head direction. The tree-trunk trials are divided into East-wall (blue) and West-wall (red) epochs, with (viewed from the camera’s perspective) 90 degrees being “up” and zero degrees being “right” etc. Firing rates at the peak direction are shown in black for the baseline and red and blue for West and East values, respectively. Note that the firing directions are 180 degrees opposed. (B) Calculation of separation between East and West polar plots – the plots from each wall are cross-correlated, with the peak being the amount of rotation that produces the highest correlation. This is close to 180 degrees for both cells. (C) Circular plot of the angular difference between firing on the two walls, for the cells that fired on both walls (all but four). Each dot represents the value calculated for one cell, based on the cross-correlation method (see Experimental Procedures and Supplementary info). The black line represents the mean direction for all cells. (D) Comparison of the actual firing directions on the West wall against the value predicted by each of the three hypothetical models (data are jittered in the x-axis for ease of visualization). The dotted line shows zero deviation: the mosaic predictions lay closest to this value, and were significantly different from the global prediction (** = p < 0.01).

**Figure 4.**
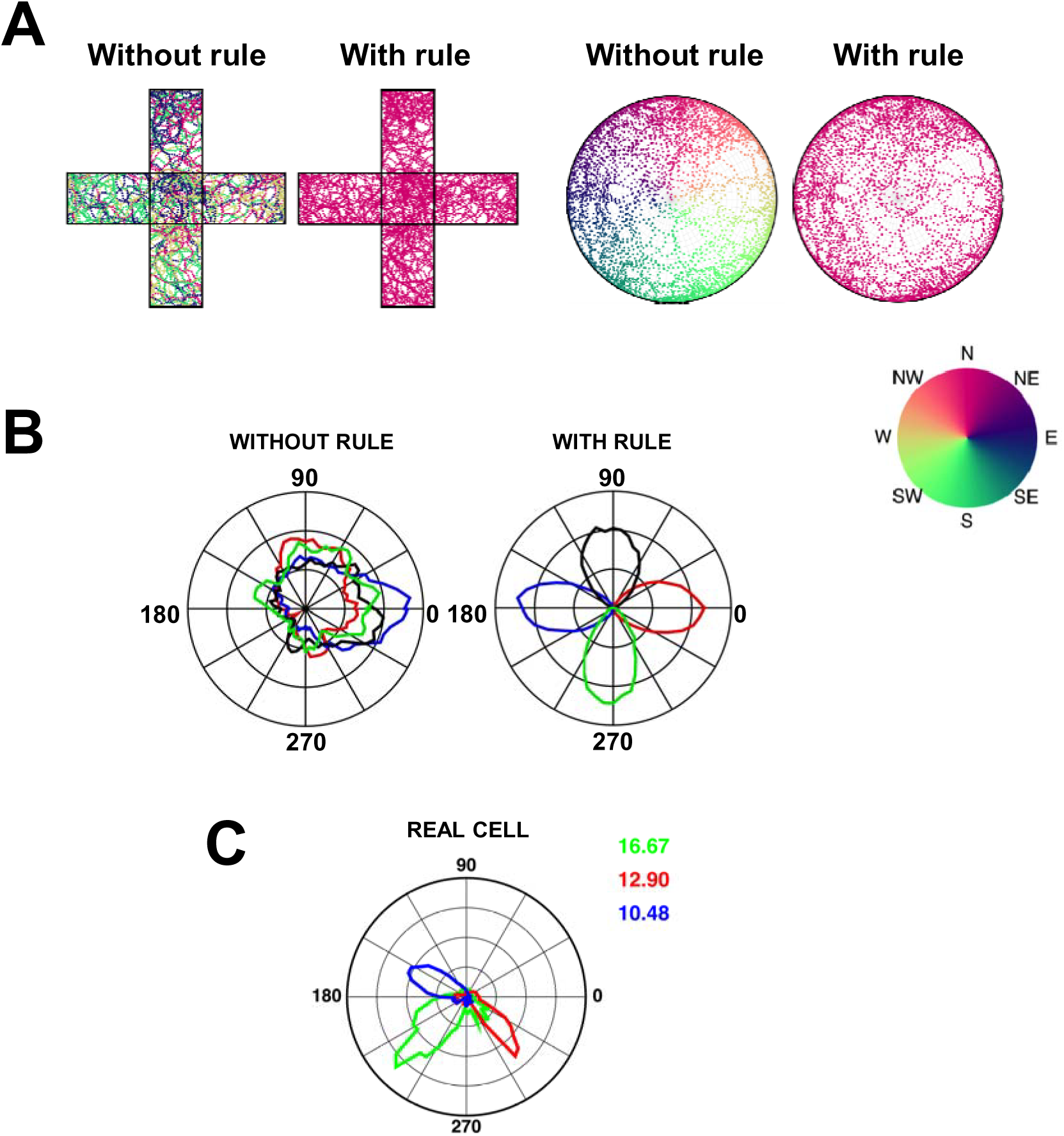
Simulating the effect of the gravity-vector rule on the consistency of head direction cell firing on three-dimensional surfaces. The two left panels illustrate the pattern as seen from above on the unfolded cuboid apparatus, and the two right panels are the pattern seen from above on a hemisphere. Positions of a simulated exploring rat were sampled every 100 ms: each dot represents the current state of the “North” head direction cell at that point, colored according to the direction it would be firing (see color map) if the simulated rat were to move directly (i.e., without rotation) to the uppermost, horizontal surface of the apparatus. Without operation of the gravity rule, firing would be inconsistent, as shown by the variable colors on each surface. With the rule, the “North” cell would always fire when the rat faced North. The difference in error pattern between the two apparati (intermingled errors for the cuboid, smoothly transitioning errors for the hemisphere) is due to the fact that the errors that accumulate for rotations about gravity in one direction are corrected again by rotations in the other direction, except for the special case when the rat moves to the top of the apparatus directly without rotation – this rarely if ever happens on the hemisphere and always happens on the cuboid. (B) Polar plots showing firing profile of network cell 250 in the camera frame of reference for each wall for model-generated data on the simulated tree-trunk maze with (right) and without (left) the gravity rule. Firing rate data were averaged across 6 degree bins of directional heading within the current wall. Data for the East wall is plotted in blue, the West wall in red, the North wall in black, and the South wall in green. Without operation of the gravity rule (left), there is no consistent relationship between cell firing and a given heading direction within the current wall. With the gravity rule (right), firing is strongly related to heading direction within the current wall. Similarly to Fig. 3, preferred firing directions on opposing walls are separated by 180 degrees. Additionally, preferred firing directions on adjacent walls are separated by 90 degrees. (C) One cell was also recorded in the real experiment on the South wall: as expected, its firing was 90 degrees to the left of the East wall and 90 degrees to the right of the West wall, conforming to the rotation around the respective corners.

Peak firing rates were significantly higher on baseline trials than on climbing trials (East and West walls combined, except for the four one-wall cells in which just the active-wall values were used), averaging 27.43 +/‐ 5.25 Hz on the flat and 20.40 +/‐ 3.50 Hz on the walls [ t(18) = 2.32, p = 0.03]. General directionality of firing, based on Rayleigh vector scores (see Supplementary Information) averaged 0.75 +/‐0.01 in baseline trials and 0.71 +/‐ 0.02 in climbing trials; although slightly lower on the walls, these values did not significantly differ [t(18) = 1.39, p = 0.18]. This indicates that HD cell firing is equally as well modulated by yaw rotations during movement on the vertical plane as on the horizontal.

We then looked at the actual directions of firing. The firing of two example cells is shown in Fig. 3A, in which it can be seen that firing directions in each local, wall-anchored reference frame were 180 degrees apart on the East and West walls, a pattern that was observed in all the cells from all three animals. Firing direction angular separations averaged 178.13 +/‐ 9.75 degrees (Fig. 3C); a circular V-test confirmed that this was significantly concentrated around 180 degrees [V(1,14) = 14.61, p < 0.001]. We then compared the firing directions on the West wall with predictions made by transforming the East-wall firing direction according to each of the three models (Supp. Fig. 5 and Fig. 3D). The data are clustered around 180 degrees for the local model, as expected (since the local model predicts zero rotation but we have already seen that the data were clustered at 180 degrees). Neither the global nor mosaic-model deviations were different from zero – however, the global and mosaic models differed significantly [t(14) = 2.66, p < 0.01], indicating that despite their expected overlap (since global and mosaic predictions are similar for some firing directions) the mosaic model was a better predictor of the data than the global model. The large spread of the global-prediction deviations is accounted for by the fact that predicted rotations vary depending on the original firing direction, being 180 degrees for those near the horizontal axis and zero for those near the vertical (Fig. 2 B-D).

For one trial, one of the cameras was moved and an additional recording was made on the South wall – this showed a firing direction intermediate between the other two walls, lying to the left (counter-clockwise) of the East wall pattern and to the right (clockwise) of the West wall pattern (Fig. 4B).

The data above suggest a dual-rotation “mosaic” rule that combines two updating rules: one for rotations around the D-V axis, and one for rotations of that axis around the Earth-vertical axis. To test the robustness of the rule we modelled it using a simulated ring attractor (Skaggs et al. 1995; Zhang 1996; Redish et al. 1996), on both a cuboid and a hemisphere.

### Modelling

The mosaic rule is graphically depicted in Supp. Fig. 6 for an imaginary rat walking on a hemisphere. This figure shows how under these conditions, each unit of movement over the surface incurs both a rotation around the D-V axis of the rat and also a rotation of the D-V axis around the vertical axis.

We modelled this dual-rotation-rule proposition in a ring attractor, which is the classic network used to model HD cells (Skaggs et al. 1995; Zhang 1996; Redish et al. 1996). A simulated rat explored two three-dimensional environments: the tree-trunk maze and a hemisphere. The model was initialized with the imaginary rat facing “North” and the activity in the ring centered over the North head direction cell; it was then updated every 100 ms. Fig. 4 shows the activity of the head direction network after some minutes of exploration. Using only a local rule, head direction cell firing progressively deviates in both of the three-dimensional environments. On the tree-trunk, errors occur abruptly when the rat crosses a corner, executing a sudden rotation around the vertical axis. On the hemisphere the errors accumulate gradually, because each step rotates the animal incrementally around the vertical axis, except in the case where the animal heads straight towards the upper pole. In the presence of the mosaic rule, by contrast activity is always correct in both environments. Note that the rule would work for both convex surfaces, when the D-V axis slopes away from the vertical axis, and concave surfaces where it slopes towards it (not shown here).

Fig. 4B shows model behavior at the single-cell level for an example cell. In the absence of the mosaic rule, cell firing does not have a consistent relationship to heading in the current wall’s camera reference frame. In contrast, during simulations incorporating the mosaic rule, the cell shows a clear preferred firing direction on a given wall, with transitions between walls causing a 90 degree rotation of preferred firing direction. Thus, the model has replicated experimental findings on both the East/West walls and the South wall.

## Discussion

We have shown that the activity of head direction cells as animals move over a three-dimensional surface may be organized by a “mosaic” updating rule, in which firing directions are updated not just by yaw rotations about the animal’s D-V axis but also rotations of the D-V axis around the vertical axis. We called this rule “mosaic” because it allows generation of multiple planar local directional reference frames which can be linked together in global 3D space without inducing conflicts (Figs 1E and 4A).

The issue of conflicts arising in directional reference frames for animals moving in 3D was first raised by Knierim et al (Knierim et al. 2000), who found evidence that place cells recorded in microgravity are somehow able to avoid these conflicts. The mosaic rule offers a way to do this, because it allows a simple, planar updating rule to operate locally, but also takes into account the spatial relation of differently oriented planes to each other. Our data, showing appropriate rotation of HD cell firing for rats moving on opposing vertical walls, provide experimental support for the mosaic rule. The mosaic rule also accounts for the behavior of four cells that switched off on one of the walls: this can be explained by noting that the mosaic rule would predict firing in a direction the rats did not sample often. With modelling we then confirmed that the mosaic updating rule allows activity to be moved smoothly around a ring attractor [the usual architecture for modelling HD neurons (Skaggs et al. 1995; Zhang 1996; Redish et al. 1996)] without discontinuities or conflicts (note that we restricted analysis to cases where the animal remained with its dorsal surface uppermost: the case of inversion is discussed further below) and it generalizes from a cuboid to free movement over a hemisphere, and would also generalize to concave surfaces (not shown).

Our experiment builds on previous work by Stackman et al (Stackman et al. 2000) and Taube et al (Taube et al. 2012) which showed that head direction cells maintained firing directions as rats moved under their own volition from a floor to a wall, with the result that depending on the cardinal direction of the wall, a different cell would fire when the rat faced upwards on walls oriented in different directions. These authors proposed a hemi-torus model in which the cells’ activity could be accounted for by a simple rotation of the floor’s horizontal coordinate system to any angle between +/‐ 90 degrees. Our experiment extends this finding by revealing active rotation of the HD cell reference frame during transitions between differently oriented walls, invoking the need for an additional rotation-based updating rule. Preservation of firing direction between the floor and the wall, as in Stackman et al (2000) and Taube et al (2012) can be accounted for by the fact that this rotation, around a horizontal axis, has no component around the Earth-vertical axis and thus invokes no HD rotation.

Our analyses, both experimental and theoretical, were restricted to movement of an upright animal: what if the animal becomes inverted? We avoided modelling this scenario because on inversion, the D-V axis of the rat, which is central to the mosaic rule, reverses through zero (in which azimuth is undefined) from positive (dorsal up) to negative (ventral up), which would have necessitated a sudden jump of the network state from one “side” of the ring attractor to the other. This singularity in the mosaic model offers a potential explanation for previous observations that in rats, a pitch (head-over-heels) rotation to inversion causes HD firing to degrade (Calton 2005). In bats, azimuth reverses instead, permitting the network state to remain unaltered (i.e., cells keep firing (Finkelstein et al. 2015); see (Jeffery et al. 2015) for more detailed discussion of this issue).

The mosaic rule combines egocentric and allocentric reference frames – egocentric from rotation of the animal around its own D-V axis, and allocentric from rotation of the D-V axis around gravity – and thus allows a directional reference frame to be constantly consistent in global space. Our experiments were conducted in rats: in bats it appears that head direction cells have tuning curves specific to pitch angle (unlike rats in which pitch-sensitive cells appear modulated by, rather than tuned to, pitch (Stackman & Taube 1998)), and many cells show conjunctive encoding suggestive of a more volumetric encoding scheme than the planar one evident here (Finkelstein et al. 2015). However, the population of azimuth-specific cells far exceeds those specific to roll or pitch, suggesting that even in these animals the encoding has a planar bias, although the dominant plane may be the Earth-horizontal rather than the locomotor surface as appears to be the case with rats.

Which brain regions might be involved in detection of rotation around the Earth-vertical axis? Horizontal rotation is detected with the labyrinthine semicircular canals(Brown 1874; Angelaki & Cullen 2008), but gravity, which is a form of linear acceleration, is detected by the saccule in the otolith organs. Angelaki and colleagues have shown that neurons in the vestibular nuclei that receive otolith afferents have response properties that would contribute determination of the gravity vector relative to off-axis rotation of the head (Angelaki 1992a; Angelaki 1992b). Another structure likely to be involved is the vestibulocerebellum, downstream of the vestibular nuclei, where neurons extract the gravity vector from a combination of otolith and semi-circular canal signals (Angelaki & Hess 1995). There are abundant projections from here to the head direction circuit (see Shinder and Taube (Shinder & Taube 2010) for review).

Our investigations only pertain to movement over a surface – different issues arise for animals that can move freely through a volumetric space, as bats, birds and fish do. Experiments in bats suggest that head direction cells also encode pitch (Finkelstein et al. 2015), and many cells show conjunctive encoding suggestive of a more volumetric encoding scheme than the planar one evident here. However, even in bats the population of azimuth-specific cells far exceeds those specific to roll or pitch, suggesting that even in these animals the encoding has a planar bias, although the dominant plane may be the Earth-horizontal rather than the locomotor surface, as with rats.

In summary, the mosaic rule allows a multi-planar, mosaic-like representation of heading over complex terrain, in which local representations of heading are linked by their relations with respect to the gravity-defined vertical axis. In this way, a stable sense of direction can be maintained at any point, without errors or conflicts, allowing animals to remain oriented as they navigate over complex terrain.

## Experimental procedures

### Electrophysiology

Details of the experimental procedures are available in the Supplementary Information. Briefly, rats were raised in enriched housing (a large aviary) to develop climbing skills, and then implanted with microdrives (Axona Ltd, Herts, UK) each carrying four tetrodes aimed at the anterodorsal nucleus of the thalamus (ADN). Once head direction cells were identified, each animal underwent a recording sequence comprising baseline foraging trials on a horizontal 70 x 70 cm box with 50 cm high walls, followed by climbing trials on a cuboid “tree trunk maze” which had walls 80 cm high and 50 cm wide, covered with chicken-wire to aid climbing. Attached to the base of the South wall was a starting box measuring 50 x 30cm and with 20 cm high walls on the three exposed sides, from which rats would start all trials. Two cameras were positioned facing the East and West walls, in order to track position and heading. In one of the trials for one animal, one of the cameras was moved in order to track climbing on the South wall as well.

During climbing, unit data were collected via a DacqUSB recording system (Axona Ltd) while the position and heading of the animal were tracked by means of LEDs on the headstage. Data were analyzed offline using a cluster-cutting program (Tint, Axona Ltd) to extract spikes and their associated position and head direction. Directional firing was determined by binning the firing in 64 bins, Gaussian smoothing and determining the Rayleigh vector score (Batschelet 1981). Relative firing directions on the East vs West walls was determined by a cross-correlation procedure between the East and West wall data sets, after which the peak was extracted to determine angular separation of firing on the two walls.

### Modelling

Details of the model are provided in the Supplementary Information. The model is based on a Continuous Attractor Neural Network (CANN), a well-established architecture for modelling the head direction system. A ring of head direction cells is topologically organized with preferred firing directions, plus a Gaussian activity packet within this network to represent current head direction. The cells are connected by pre-wired fixed-weight recurrent collateral synapses, with the strength of each connection depending on the similarity of preferred firing directions. The cells are modelled as a leaky-integrator firing rate neurons (described in more detail in the Supplementary Information) in which activity is represented as an instantaneous average firing rate for each cell, with an axonal delay between one cell and the next. All head direction cells receive self-motion input reflecting both the change in heading of the rat within the current horizontal reference plane relative to the D-V axis, and rotations of this D-V axis about the earth-referenced vertical axis defined by the gravity vector. This input is a Gaussian, centered on a point determined as the previous ring location to which input was applied at the previous timestep together with an increment based on the combined amount of D-V-axis and gravity-vector rotations.

Details of the model are provided in the Supplementary Information. The model is based on a Continuous Attractor Neural Network (CANN), a well-established architecture for modelling the head direction system. A ring of head direction cells is topologically organized with preferred firing directions, plus a Gaussian activity packet within this network to represent current head direction. The cells are connected by pre-wired fixed-weight recurrent collateral synapses, with the strength of each connection depending on the similarity of preferred firing directions. The cells are modelled as a leaky-integrator firing rate neurons (described in more detail in the Supplementary Information) in which activity is represented as an instantaneous average firing rate for each cell, with an axonal delay between one cell and the next. All head direction cells receive self-motion input reflecting both the change in heading of the rat within the current horizontal reference plane relative to the D-V axis, and rotations of this D-V axis about the earth-referenced vertical axis defined by the gravity vector. This input is a Gaussian, centered on a point determined as the previous ring location to which input was applied at the previous timestep together with an increment based on the combined amount of D-V-axis and gravity-vector rotations.

Custom-built MATLAB scripts were used to generate a random walk either over the surface of a cuboid or a hemisphere (the cuboid notionally having dimensions identical to the tree trunk apparatus, namely 50 × 50 × 80 cm; the hemisphere having radius 50cm), located in an imaginary allocentric reference frame. Every 100 ms in model time the simulated rat took a “step” of length 2.5 cm, the direction of which was selected from a Gaussian probability distribution centered ahead of the rat. For both the cuboid and hemisphere, boundary conditions were imposed when the rat neared the bottom of the apparatus, such that the Gaussian probability distribution determining step direction was shifted away from the bottom edge. Trials were 600 seconds long, generating 6000 pairs of position points, each pair corresponding to the beginning and end of a step. If a rotation about the D-V axis, *or* of the D-V axis about the vertical axis, occurred during a step, self-motion input was updated so as to shift the head direction firing packet location accordingly. The azimuth correlate of the new head direction cell firing direction was then calculated. At each time point the rat’s current head direction was determined by effectively transporting the simulated rat by a direct path to the uppermost point of the maze, thus restoring it to horizontal without rotating it. The firing direction of the “North” cell was determined at this point, and compared against actual North to generate an error estimate. This value was then plotted on the corresponding point on the sphere, using a color code to indicate error. On each wall, current heading was recoded into a head direction in the current camera frame of reference by taking the angle between current heading and a reference vector pointing directly upwards. This was then related to the firing of a given example cell to determine that cell’s preferred firing direction.

## Acknowledgements

This work was supported by BBSRC (BB/J009792/1), MRC (G1100669) and Wellcome (#083540) grants to KJ, and a BBSRC CASE studentship (BB/F015968/1) to JW in collaboration with Axona Ltd. KJ is a non-shareholding director of Axona Ltd.

